# A DNase T6SS effector requires its MIX domain for secretion

**DOI:** 10.1101/2022.05.04.489851

**Authors:** Chaya Mushka Fridman, Biswanath Jana, Rotem Ben-Yaakov, Eran Bosis, Dor Salomon

**Author notes:** Address correspondence to Dor Salomon. Chaya Mushka Fridman and Biswanath Jana contributed equally to this work. Author order was determined on the basis of earlier involvement in the project. Present address: Rotem Ben-Yaakov, TransAlgae, Rehovot, Israel.

## Abstract

Gram-negative bacteria often employ the type VI secretion system (T6SS) to deliver diverse cocktails of antibacterial effectors into rival bacteria. In many cases, even when the identity of the delivered effectors is known, their toxic activity and mechanism of secretion are not. Here, we investigate VPA1263, a *Vibrio parahaemolyticus* T6SS effector that belongs to a widespread class of polymorphic effectors containing a MIX domain. We reveal a C-terminal DNase toxin domain belonging to the HNH nuclease superfamily, and we show that it mediates the antibacterial toxicity of this effector during bacterial competition. Furthermore, we demonstrate that the VPA1263 MIX domain is necessary for T6SS-mediated secretion and intoxication of recipient bacteria. These results are the first indication of a functional role for MIX domains in T6SS secretion.

**IMPORTANCE:** Specialized protein delivery systems are used during bacterial competition to deploy cocktails of toxins that target conserved cellular components. Although numerous toxins have been revealed, the activity of many remains unknown. In this study, we investigated such a toxin from the pathogen *Vibrio parahaemolyticus*. Our findings indicated that the toxin employs a DNase domain to intoxicate competitors. We also showed that a domain used as a marker for secreted toxins is required for secretion of the toxin via a type VI secretion system.

## INTRODUCTION

Bacterial polymorphic toxins are modular proteins delivered by diverse secretion systems to mediate antibacterial or anti-eukaryotic activities (1, 2). They often share an N-terminal domain fused to diverse C-terminal toxin domains. The N-terminal domain, which can be used to classify these toxins, may determine which secretion system the effectors will be secreted through, e.g., the type V, VI, or VII secretion systems (T5/6/7SS, respectively) (1, 3–6).

Many polymorphic toxins are secreted via T6SS, a contractile injection system widespread in Gram-negative bacteria (7, 8). The toxins, called effectors, decorate a secreted arrow-like structure comprising an inner tube, made of stacked Hcp hexamers, and a capping spike containing a VgrG trimer and a PAAR repeat-containing protein (9–12); the effector-decorated arrow is propelled outside of the cell by a contracting sheath structure that engulfs the inner tube (13). Effectors can be either specialized – Hcp, VgrG, or PAAR proteins containing additional toxin domains at their C-termini – or cargo effectors, which are proteins that non-covalently bind to Hcp, VgrG, or PAAR with or without the aid of an adaptor protein or a co-effector (14–21). To date, three classes of polymorphic T6SS cargo effectors containing MIX (3), FIX (22), or Rhs (23) domains have been characterized; the Rhs domain is not restricted to T6SS effectors (23).

Proteins belonging to the MIX-effector class are secreted via T6SS (3, 24–26). They contain a predominantly N-terminal MIX domain fused to known or predicted anti-eukaryotic or antibacterial C-terminal toxin domains; the latter precede a gene encoding a cognate immunity protein that prevents self or kin-intoxication (4, 27, 28). MIX domains can be divided into five clans that differ in their sequence conservation (3, 29). Notably, members of the MIX V clan are common in bacteria of the *Vibrionaceae* family (3, 29) and were suggested to be horizontally shared via mobile genetic elements (25). It remains unknown whether MIX plays a role in T6SS-mediated secretion.

In previous work, we identified the MIX-effector VPA1263 as an antibacterial effector delivered by *V. parahaemolyticus* RIMD 2210633 T6SS1, and its downstream encoded Vti2 as the cognate immunity protein (3) (**Fig. 1A**). VPA1263 belongs to the MIX V clan. It is encoded on *V. parahaemolyticus* island-6 (VpaI-6), a predicted mobile genomic island that is present in a subset of *V. parahaemolyticus* genomes (30). In this work, we aimed to investigate the toxic activity and the secretion mechanism of VPA1263. We found that VPA1263 contains a C-terminal toxin domain belonging to the HNH nuclease superfamily, and showed that this domain functions as a DNase. Furthermore, we found that the MIX domain is required for secretion of VPA1263 via the T6SS, providing the first experimental validation of the hypothesis that MIX domains play a role in T6SS effector secretion.

**Figure 1.**
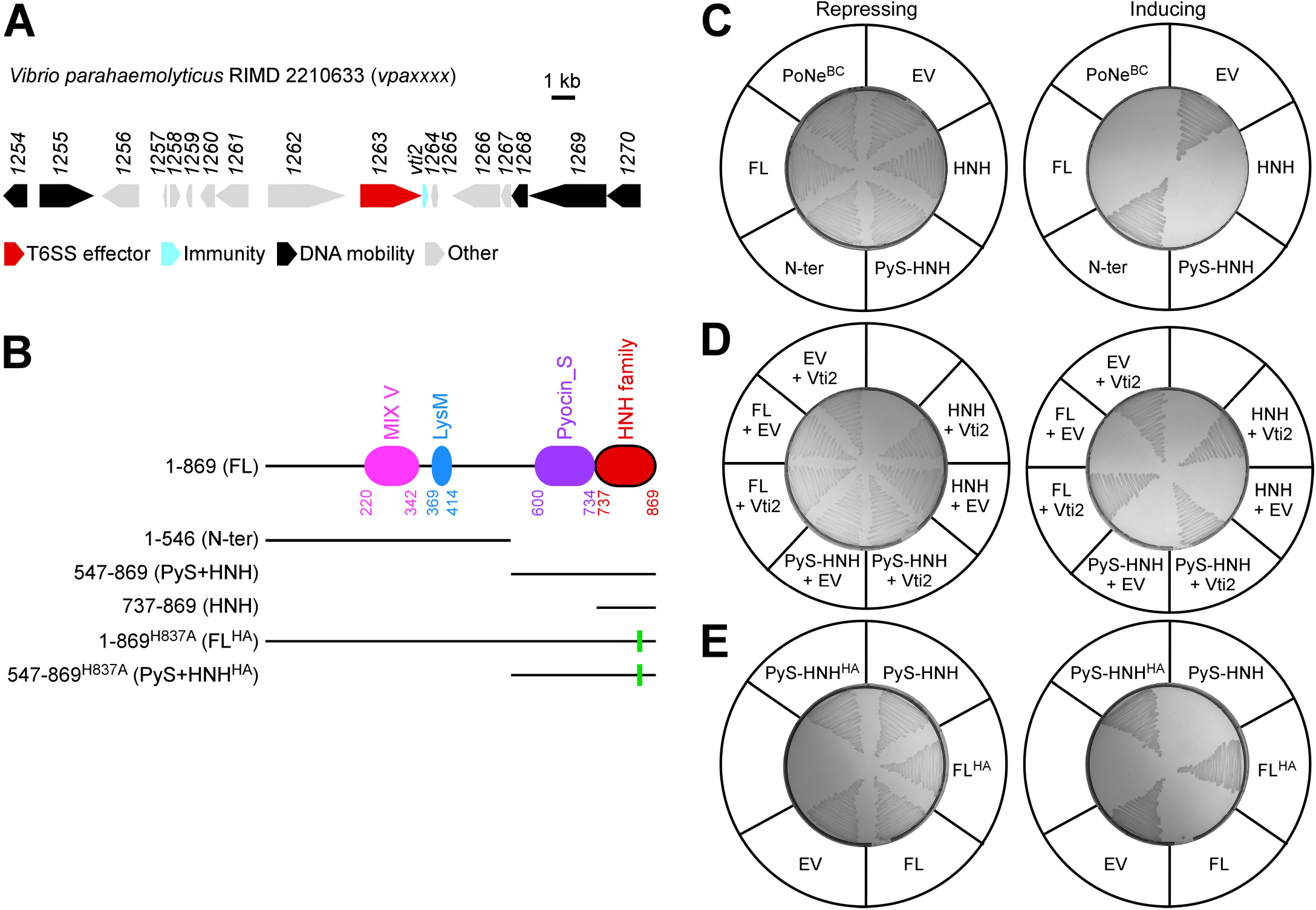
A C-terminal domain mediates VPA1263’s antibacterial toxicity. **(A)** The *vpa1263* gene neighborhood (genome accession number BA000032.2). Genes are denoted by arrows indicating the direction of transcription. Locus tags (*vpaxxxx*) and unannotated gene names (i.e., *vti2*) are shown above. **(B)** Schematic representation of the domains identified in VPA1263 and of truncated and mutated variants examined in this figure. Green rectangles denote the position of the H837A mutation. **(C-E)** Toxicity of C-terminal Myc-His-tagged VPA1263 variants expressed from an arabinose-inducible expression plasmid in *E. coli* MG1655 streaked onto repressing (glucose) or inducing (arabinose) agar plates. In **D**, a second arabinose-inducible plasmid was used to co-express Vti2. EV, empty plasmid; PoNe^BC^, a PoNe domain-containing DNase from *Bacillus cereus* (BC3021). The results shown represent at least three independent experiments.

## RESULTS

### VPA1263 contains a C-terminal HNH nuclease-like toxin domain

Analysis of the amino acid sequence of VPA1263 using the NCBI conserved domain database (31) revealed three known domains: a MIX domain (3), followed by a peptidoglycan-binding LysM domain (32), and a Pyocin_S domain (33) (**Fig. 1B**). Hidden Markov modeling using HHPred (34) revealed another region at the C-terminus of VPA1263 (amino acids 737-869) that is similar to the toxin domain found in members of the HNH nuclease superfamily, such as antibacterial S-type pyocins and colicins (35–40). An iterative search using a hidden Markov Model profile against the UniProt protein database indicated that the Pyocin_S and putative HNH nuclease domains located at the C-terminus of VPA1263 are also found together at the C-termini of specialized T6SS effectors containing PAAR, Hcp, or VgrG (**Supplementary Fig. S1A**); they were also found together in colicins such as colicin E9 and E7, although sometimes they were separated by a receptor binding domain. Interestingly, a similar HNH nuclease-like domain was also found in effectors containing N-terminal domains such as LXG (5) and WXG100 (41), which are associated with T7SS, but without an accompanying Pyocin_S domain. A conservation logo generated by aligning the amino acid sequences of the VPA1263 C-terminal domain (737-869) and homologous domains found in known or predicted secreted toxins confirmed the presence of a conserved His-Gln-His motif (42) (**Supplementary Fig. S1B**). These results led us to hypothesize that VPA1263 contains a C-terminal HNH nuclease domain that can mediate its antibacterial toxicity.

To determine the minimal region sufficient for VPA1263-mediated antibacterial toxicity, we ectopically expressed VPA1263 or its truncated versions (**Fig. 1B**) in *E. coli*. As shown in **Fig. 1C**, the C-terminal end, corresponding to the predicted HNH nuclease domain (amino acids 737-869; HNH), was necessary and sufficient to intoxicate *E. coli*. Notably, the expression of all VPA1263 versions, except for the toxic HNH domain, was detected in immunoblots (**Supplementary Fig. S2A**). Co-expression of the cognate immunity protein, Vti2 (3), rescued *E. coli* from the toxicity mediated by either the full-length VPA1263 or the C-terminal HNH nuclease domain (**Fig. 1D**), confirming that the toxicity mediated by the C-terminal HNH nuclease domain resulted from the same activity that is used by the full-length VPA1263. In addition, replacing the conserved histidine 837 with alanine (FL^HA^ or PyS-HNH^HA^) abrogated VPA1263-mediated toxicity (**Fig. 1E**), further supporting the role of the HNH nuclease domain in VPA1263-mediated antibacterial toxicity. Notably, the expression of the mutated proteins was confirmed in immunoblots (**Supplementary Fig. S2B**). Taken together, our results suggest that VPA1263 exerts its antibacterial toxicity via a C-terminal HNH nuclease domain.

### VPA1263 is a DNase

Since HNH nucleases target DNA, we hypothesized that VPA1263 is a DNase. To test this hypothesis, we first set out to determine whether the C-terminal domain of VPA1263 can cleave DNA in vitro and in vivo. Since we were unable to purify the minimal toxin domain (amino acids 737-869), possibly due to low expression levels, we purified a truncated version encompassing both the C-terminal Pyocin_S and HNH nuclease domains (amino acids 547-869; PyS-HNH) (**Supplementary Fig. S3A**). As expected, this toxic region was sufficient to cleave purified *E. coli* genomic DNA in vitro in the presence of MgCl_2_, similar to DNase I, which was used as a control (**Fig. 2A**). In contrast, a mutated version, in which histidine 837 was replaced with alanine (PyS-HNH^HA^), did not cleave the DNA. Furthermore, we were only able to isolate small amounts of genomic DNA from *E. coli* cultures expressing an arabinose-inducible PyS-HNH or the *Bacillus cereus* DNase, BC3021 (PoNe^BC^; (22)) (**Fig. 2B**), indicating that VPA1263 also cleaves DNA in vivo.

**Figure 2.**
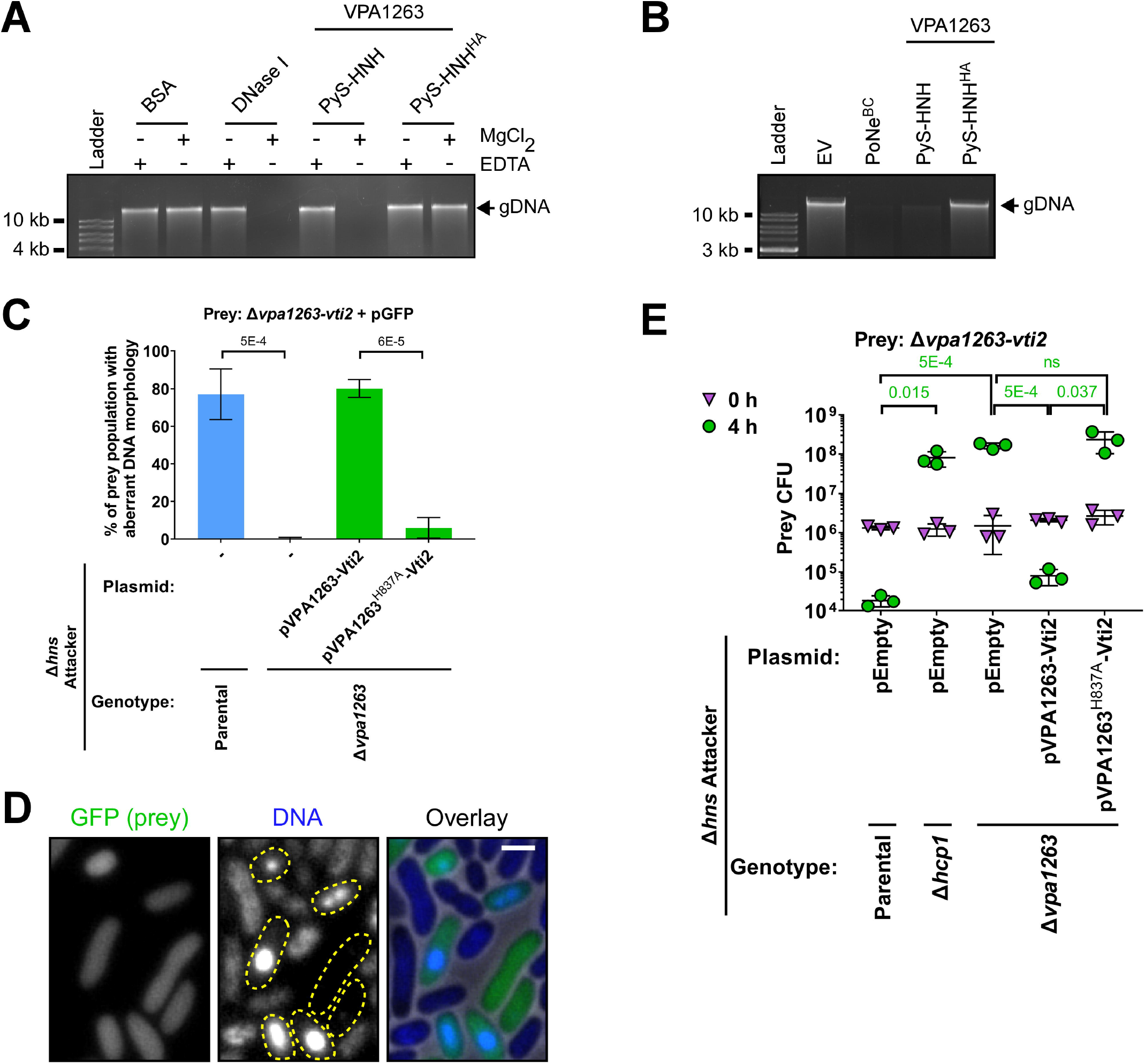
VPA1263 is a DNase. (A) In vitro DNase activity. Purified *E. coli* genomic DNA (gDNA) was co-incubated with the indicated purified protein in the presence (+) or absence (−) of MgCl_2_ or EDTA for 5 min at 30°C. The results shown represent two independent experiments. **(B)** In vivo DNase activity. The integrity of gDNA was determined after its isolation from *E. coli* MG1655 cells in which the indicated proteins were expressed from arabinose-inducible expression plasmids. The results shown represent at least three independent experiments. EV, empty plasmid; PyS-HNH, amino acids 547-869 of VPA1263; PyS-HNH^HA^, PyS-HNH with a H837A mutation; PoNe^BC^, a PoNe domain-containing DNase from *Bacillus cereus* (BC3021). **(C)** Quantification of the percentage in the population of *V. parahaemolyticus* RIMD 2210633 Δ*vpa1263*-*vti2* prey cells showing aberrant DNA morphology (no detectable DNA or DNA foci) after 3 h of competition against the indicated attacker strains. DNA was visualized by Hoechst staining. The results represent the average ± SD of three independent experiments. Statistical significance between samples by an unpaired two-tailed Student’s *t*-test is denoted above. In each experiment, at least 100 prey cells were randomly selected and manually evaluated per treatment. **(D)** Sample fluorescence microscope images of prey cells after 3 h competition against a Δ*hns* attacker strain, as described in **C**. Dashed yellow shapes in the DNA channel encircle prey cells detected in the GFP channel. Bar = 1 μm. **(E)** Viability counts of the indicated prey strain before (0 h) and after (4 h) co-incubation with the indicated *V. parahaemolyticus* RIMD 2210633 Δ*hns* derivative attacker strain on MLB agar plates supplemented with L-arabinose at 30°C. Prey strains contain an empty plasmid that provides a selection marker. Data are shown as the mean ± SD, n = 3 technical replicates. Statistical significance between samples at the 4 h timepoint by an unpaired two-tailed Student’s *t*-test is denoted above. The experiment was performed three times with similar results; results of a representative experiment are shown.

Next, we set out to determine whether VPA1263 targets prey DNA during T6SS-mediated bacterial competition. To this end, we used fluorescence microscopy to visualize the DNA in bacterial cultures after T6SS-mediated self-competition. We competed a *V. parahaemolyticus* prey strain that constitutively expresses a green fluorescent protein (GFP; used to distinguish prey from attacker cells) in which we deleted *vpa1263* and *vti2* (to specifically sensitize it to VPA1263-mediated toxicity) against attacker strains with a constitutively active T6SS1 deleted for the T6SS1 repressor *hns* to maximize T6SS1 activity (43), with or without a deletion in *vpa1263* (Δ*hns*/Δ*vpa1263* and Δ*hns*, respectively). Notably, these attacker strains exhibited similar growth rates (**Supplementary Fig. S3B**), and the Δ*hns*/Δ*vpa1263* strain retained the ability to intoxicate *Vibrio natriegens* prey in competition, indicating that it was able to deliver other T6SS1 effectors (3); thus, T6SS1 remained functional even though this attacker strain was unable to intoxicate a Δ*vpa1263-vti2* prey strain (**Supplementary Fig. S3C**). As shown in **Fig. 2C,D** and in **Supplementary Fig. S4**, VPA1263-sensitive, GFP-expressing prey cells devoid of DNA or containing DNA foci, which are probably regions of stress-induced DNA condensation (44), were only detected after competition against the VPA1263^+^ attacker strain (Δ*hns*). Moreover, a plasmid for expression of VPA1263 and its cognate immunity protein, Vti2, (pVPA1263-Vti2) complemented this phenotype in a Δ*hns*/Δ*vpa1263* attacker strain background, whereas a similar plasmid for expression of the catalytically inactive mutant, VPA1263^H837A^ (pVPA1263^H837A^-Vti2), did not (**Fig. 2C**). Taken together with the inability of a plasmid-encoded catalytically inactive mutant to complement VPA1263-mediated toxicity of a Δ*hns*/Δ*vpa1263* attacker strain during bacterial competition (**Fig. 2E**), these results support the conclusion that VPA1263 is a T6SS1 effector that exerts its toxicity during bacterial competition via its DNase activity.

### The MIX domain is necessary for T6SS-mediated secretion of VPA1263

After determining that its toxicity is mediated by DNase activity, we next sought to investigate the role of VPA1263’s MIX domain. Since MIX domains are found at N-termini of diverse polymorphic toxins that are secreted by T6SS (3), we hypothesized that MIX plays a role in T6SS-mediated secretion. To test this, we monitored the T6SS1-mediated secretion of C-terminal Myc-His tagged VPA1263 variants expressed from a plasmid in a *V. parahaemolyticus* Δ*hns* strain with a constitutively active T6SS (T6SS1^+^), or in a Δ*hns*/Δ*hcp1* mutant with an inactive T6SS1 (T6SS1^−^). Notably, to avoid self-intoxication of strains over-expressing VPA1263 variants containing the HNH nuclease domain, we used the catalytically inactive mutant H837A (**Fig. 1E**). The full-length protein (FL^HA^) was secreted into the growth medium in a T6SS1-dependent manner, confirming previous comparative proteomics results (3) (**Fig. 3A**). An N-terminal region, including the MIX and LysM domains (N-ter), retained the ability to secrete via T6SS1, whereas the C-terminal region, containing the Pyocin_S and the HNH nuclease toxin domains (PyS-HNH^HA^), did not (**Fig. 3A**), indicating that the information required for T6SS1-mediated secretion is found at the N-terminal region. Remarkably, deletion of the region encoding amino acids 226-328 (ΔMIX^HA^), encompassing most of the MIX domain, resulted in loss of T6SS1-mediated secretion. This result suggests that MIX is required for T6SS1-mediated secretion of VPA1263. In support of this conclusion, replacing residues belonging to the invariant GxxY motif in the MIX domain (3) to alanine, i.e., glycine 247 and tyrosine 250 (FL^GA/HA^ and FL^YA/HA^, respectively), also abolished VPA1263’s secretion via T6SS1. Notably, we validated that T6SS1 was functional in the strains expressing the VPA1263 variants that were not secreted by detecting the T6SS1-mediated secretion of the hallmark T6SS protein, VgrG1 (**Fig. 3A**). These results provide the first experimental indication that MIX is required for T6SS-mediated secretion of an effector. This conclusion was further supported by bacterial competition assays, in which VPA1263 with a mutation in the invariant glycine of the MIX domain (G247) was unable to complement the loss of prey intoxication by a Δ*hns*/Δ*vpa1263* attacker (**Fig. 3B**). Attempts to determine whether the MIX domain is sufficient to mediate secretion via T6SS were inconclusive, due to the low expression level of VPA1263 truncations containing only the MIX domain region.

**Figure 3.**
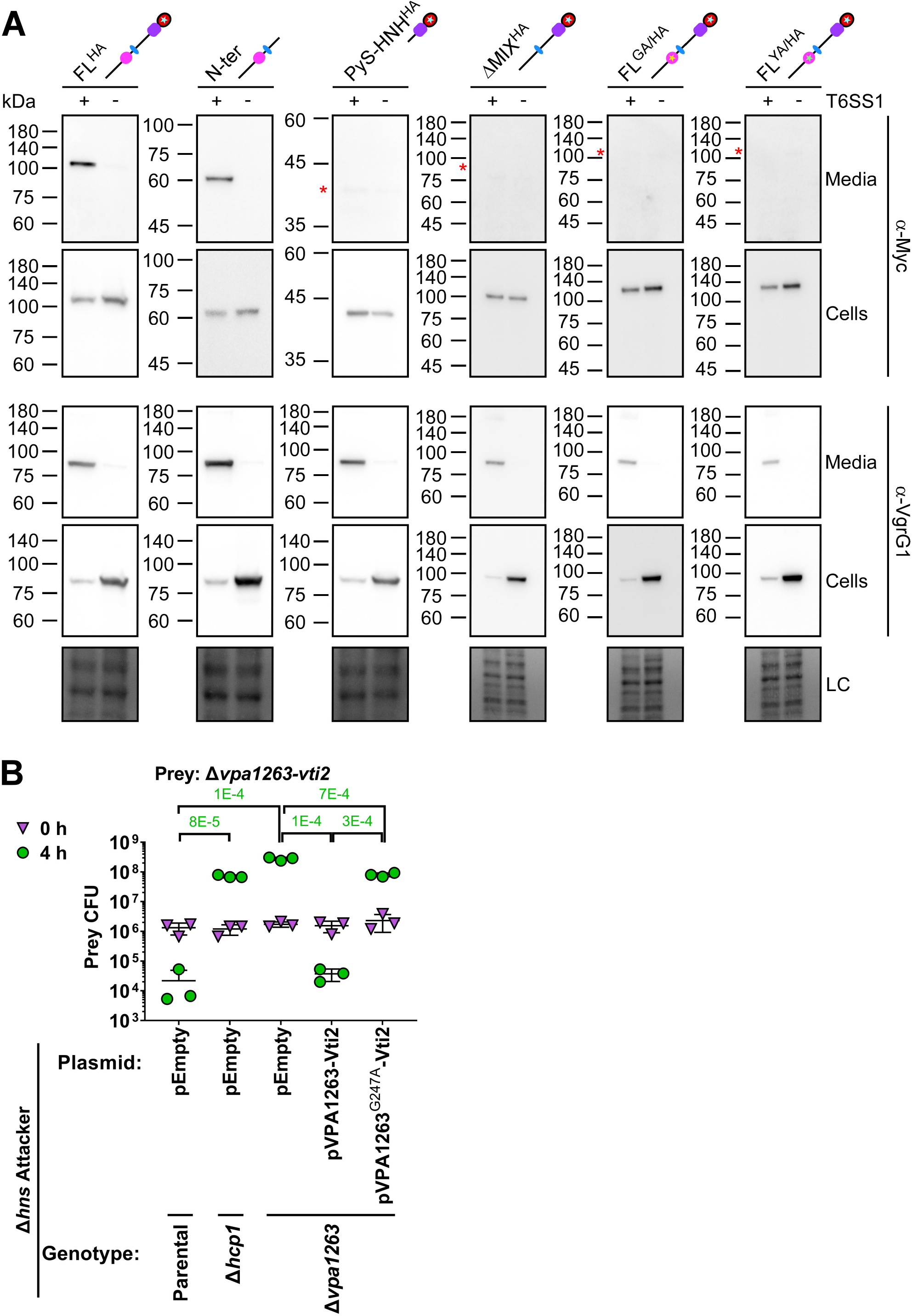
MIX is required for secretion of VPA1263 via T6SS1. (A) Expression (cells) and secretion (media) of the indicated C-terminal Myc-His-tagged VPA1263 variants from *V. parahaemolyticus* RIMD 2210633 Δ*hns* (T6SS1^+^) or Δ*hns*Δ*hcp1* (T6SS1^−^). Proteins were expressed from an arabinose-inducible plasmid, and samples were resolved on SDS-PAGE. VPA1263 variants and the endogenous VgrG1 were visualized using specific antibodies against Myc and VgrG1, respectively. FL^HA^, VPA1263 with an H387A mutation; N-ter, amino acids 1-546 of VPA1264; PyS-HNH, amino acids 547-869 of VPA1263; PyS-HNH^HA^, PyS-HNH with a H837A mutation; FL^GA/HA^, FL^HA^ with a G247A mutation; FL^YA/HA^, FL^HA^ with a Y250A mutation. Schematic representations of the expressed VPA1263 variants are shown above; denoted domains and colors correspond to Figure 1B; stars denote point mutations in the expressed variant. Red asterisks denote the expected size of the indicated proteins in the media fractions. Loading control (LC) is shown for total protein lysates. **(B)** Viability counts of the indicated prey strain before (0 h) and after (4 h) co-incubation with the indicated *V. parahaemolyticus* RIMD 2210633 Δ*hns* derivative attacker strain on MLB agar plates supplemented with L-arabinose at 30°C. Prey strains contain an empty plasmid that provides a selection marker. Data are shown as the mean ± SD, n = 3 technical replicates. Statistical significance between samples at the 4 h timepoint by an unpaired two-tailed Student’s *t*-test is denoted above. The experiment was performed three times with similar results; results of a representative experiment are shown.

## DISCUSSION

In this work, we investigated the T6SS MIX-effector VPA1263; we identified the role of its C-terminal toxin domain and of its MIX domain. Our results revealed that VPA1263 is a DNase toxin that requires its MIX domain for T6SS-mediated secretion.

Using a combination of toxicity assays, in vivo and in vitro biochemical assays, and fluorescence microscopy, we confirmed our computational prediction that VPA1263 has a C-terminal DNase domain that mediates its toxicity during bacterial competition. The identified toxin belongs to the widespread and diverse HNH nuclease superfamily (35–40); homologs are found in various known and predicted bacterial toxins. Interestingly, in some instances this toxin domain is preceded by a Pyocin_S domain, as is the case in VPA1263. Pyocin_S was recently suggested to mediate the transport of DNase toxins across the inner-membrane, from the periplasm to the cytoplasm (45). This activity was shown to be mediated by specific inner-membrane proteins that serve as receptors. It will be interesting to determine whether VPA1263 and other Pyocin_S-containing T6SS effectors require this domain for transport into the cytoplasm to mediate toxicity, since it remains unclear whether T6SS effectors are delivered directly into the recipient cell cytoplasm, periplasm, or randomly into either compartment (46–48). Notably, we recently reported that VPA1263 selectively intoxicates bacterial strains when delivered via an engineered T6SS in *V. natriegens* (49). VPA1263-delivering attacker strains were toxic to *Vibrio* and *Aeromonas* strains, but had no effect on the viability of *E. coli* or *Salmonella* prey (49). It is therefore tempting to speculate that differences in the potential inner-membrane receptors of the VPA1263 Pyocin_S domain are responsible for the observed selective toxicity. If true, it may represent a previously unappreciated mode of natural resistance against T6SS effectors.

Importantly, we found that MIX is required for T6SS-mediated secretion of VPA1263; even single point mutations in the invariant GxxY motif were sufficient to abolish effector secretion and effector-mediated intoxication of sensitive recipient prey bacteria during competition. While this is the first experimental evidence of a role for MIX in secretion, the mechanism by which it contributes to secretion remains unknown. It is possible that MIX contributes toward a stable or desirable effector conformation that is required for correct loading or positioning on the T6SS. Alternatively, MIX may mediate interaction with a secreted tube or spike component. We will investigate the underlying mechanism by which MIX mediates T6SS secretion in future work.

## MATERIALS AND METHODS

### Strains and Media

For a complete list of strains used in this study, see **Supplementary Table S1**. *Escherichia coli* strains were grown in 2xYT broth (1.6% wt/vol tryptone, 1% wt/vol yeast extract, and 0.5% wt/vol NaCl) or on Lysogeny Broth (LB) agar plates containing 1% wt/vol NaCl at 37°C, or at 30°C when harboring effector expression plasmids. The media were supplemented with chloramphenicol (10 μg/mL) or kanamycin (30 μg/mL) to maintain plasmids, and with 0.4% wt/vol glucose to repress protein expression from the arabinose-inducible promoter, P*bad*. To induce expression from P*bad*, L-arabinose was added to media at 0.05-0.1% (wt/vol), as indicated.

*Vibrio parahaemolyticus* RIMD 2210633 (50) and its derivative strains, as well as *V. natriegens* ATCC 14048 were grown in MLB media (LB media containing 3% wt/vol NaCl) or on marine minimal media (MMM) agar plates (1.5% wt/vol agar, 2% wt/vol NaCl, 0.4% wt/vol galactose, 5 mM MgSO_4_, 7 mM K_2_SO_4_, 77 mM K_2_HPO_4_, 35 mM KH_2_PO_4_, and 2 mM NH_4_Cl) at 30°C. When vibrios contained a plasmid, the media were supplemented with kanamycin (250 μg/mL) or chloramphenicol (10 μg/mL) to maintain the plasmid. To induce expression from P*bad*, L-arabinose was added to media at 0.05% wt/vol.

### Plasmid construction

For a complete list of plasmids used in this study, see **Supplementary Table S2**. For protein expression, the coding sequences (CDS) of *vpa1263* (BAC62606.1) and *vti2* (Chromosome 2, positions 1344453-1344737, GenBank number BA000032.2; encoding WP_005477334.1) were amplified from genomic DNA of *V. parahaemolyticus* RIMD 2210633. Amplification products were inserted into the multiple cloning site (MCS) of pBAD^K^/Myc-His, or pBAD33.1^F^ using the Gibson assembly method (51). Plasmids were introduced into *E. coli* using electroporation. Transformants were grown on agar plates supplemented with 0.4% wt/vol glucose to repress unwanted expression from the P*bad* promotor during the subcloning steps. Plasmids were introduced into *V. parahaemolyticus* via conjugation. Transconjugants were grown on MMM agar plates supplemented with appropriate antibiotics to maintain the plasmids.

### Construction of deletion strains

The construction of pDM4-based (52) plasmids for deletion of *vpa1263*, *vpa1263*-*vti2*, and *hns* (*vp1133*) was reported previously (3, 43). In-frame deletions of *vpa1263*, *vpa1263-vti2*, and *hns* in *V. parahaemolyticus* RIMD 2210633 were performed as previously described (53). Briefly, a Cm^R^OriR6K suicide plasmid, pDM4, containing ~1 kb upstream and ~1 kb downstream of the gene to be deleted in its multiple cloning site was conjugated into *V. parahaemolyticus*, and trans-conjugants were selected on solid media plates supplemented with chloramphenicol. Then, bacteria were counter-selected on solid media plates containing 15% (wt/vol) sucrose for loss of the *sacB*-containing plasmid. Deletions were confirmed by PCR.

### Toxicity in *E. coli*

Bacterial toxicity assays were performed as previously described (21), with minor changes. Briefly, to assess the toxicity mediated by VPA1263 or its truncated versions, pBAD^K^/Myc-His plasmids encoding the indicated proteins were transformed into *E. coli* MG1655. *E. coli* transformants were streaked onto either repressing (containing 0.4% wt/vol glucose) or inducing (containing 0.05% wt/vol L-arabinose) LB agar plates supplemented with kanamycin. Chloramphenicol was also included in the media when a pBAD33.1^F^-based plasmid was used. Plates were incubated at 30°C for 16 hours, and then imaged using a Fusion FX6 imaging system (Vilber Lourmat). The experiments were performed at least three times with similar results. Results from a representative experiment are shown.

### Protein expression in *E. coli*

Protein expression in *E. coli* was performed as previously described (21), with minor changes. Briefly, to assess the expression of C-terminal Myc-tagged proteins encoded on arabinose-inducible plasmids, overnight cultures of *E. coli* MG1655 strains containing the indicated plasmids were washed twice with fresh 2xYT broth to remove residual glucose. Following normalization to OD_600_ = 0.5 in 3 mL of 2xYT broth supplemented with appropriate antibiotics, cultures were grown for 2 hours at 37°C. To induce protein expression, 0.05% (wt/vol) L-arabinose was added to the media. After having grown for 2 additional hours at 37°C, 0.5 OD_600_ units of cells were pelleted and resuspended in (2X) Tris-Glycine SDS sample buffer (Novex, Life Sciences). Samples were boiled for 5 min, and cell lysates were resolved on Mini-PROTEAN or criterion TGX Stain-Free™ precast gels (Bio-Rad). For immunoblotting, α-Myc antibodies (Santa Cruz Biotechnology, 9E10, mouse mAb; sc-40) were used at 1:1,000 dilution. Protein signals were visualized in a Fusion FX6 imaging system (Vilber Lourmat) using enhanced chemiluminescence (ECL) reagents.

### Protein purification

To purify truncated VPA1263 proteins for the in vitro DNase assays, *E. coli* BL21 (DE3) cells harboring plasmids for the arabinose-inducible expression of the indicated Myc-His-tagged VPA1263 variants and the FLAG-tagged Vti2 (the immunity protein required to antagonize the toxicity of VPA1263 variants inside bacteria) were grown for 16 hours in 2xYT media supplemented with kanamycin and chloramphenicol at 37°C. Bacterial cultures were then diluted 100-fold into fresh media and incubated at 37°C with agitation (180 rpm). When bacterial cultures reached an OD_600_ of ~ 1.0, L-arabinose was added to the media (to a final concentration of 0.1% (wt/vol)) to induce protein expression, and cultures were grown for 4 additional hours at 30°C. Cells were harvested by centrifugation at 4°C (20 min at 13,300 x *g*), followed by washing with a 0.9% (wt/vol) NaCl solution to remove residual media. Then, cells were resuspended in 3 mL lysis buffer A (20 mM Tris-Cl pH 7.5, 500 mM NaCl, 5% vol/vol glycerol, 10 mM imidazole, 0.1 mM PMSF, and 8 M urea). Urea was included in the buffer to denature the proteins and thus release Vti2 from the VPA1263 variants. Cells were disrupted using a high-pressure cell disruptor (Constant system One Shot cell disruptor, model code: MC/AA). To remove cell debris, lysates were centrifuged for 20 min at 13,300 x *g* at 4°C. Next, 250 μL of Ni-Sepharose resin (50% slurry; GE healthcare) were pre-washed with lysis buffer A and then mixed with the supernatant fractions of lysed cells containing the denatured proteins. The suspensions were incubated for one hour at 4°C with constant rotation, and then loaded onto a column. Immobilized resin was washed with 10 mL wash buffer A (20 mM Tris-Cl pH 7.5, 500 mM NaCl, 5% vol/vol glycerol, 40 mM imidazole, and 8 M urea). Bound proteins were eluted from the column using 1 mL elution buffer A (20 mM Tris-Cl pH 7.5, 500 mM NaCl, 5% vol/vol glycerol, 500 mM imidazole, and 8 M urea). The presence and purity of the eluted Myc-His-tagged VPA1263 variants were confirmed by SDS-PAGE, using stain-free gels (Bio-rad).

To refold the denatured, purified proteins before in vitro DNase activity assays, a refolding procedure was applied. The eluted proteins were dialyzed against DNase assay buffer (20 mM Tris-Cl pH 7.5, 200 mM NaCl, 5% vol/vol glycerol) and incubated for an hour at 4°C. The buffer was replaced twice to remove unwanted imidazole and urea. Then, the 1 mL suspension was concentrated to ~ 300 μL using a Spin-X UF concentrator column (Corning, 30 kDa). The purified proteins were quantified by the Bradford method using 5X Bradford reagent (Bio-rad). The procedure was carried out at 4°C.

### In vitro DNase assays

For determining in vitro DNase activity, genomic DNA isolated from *E. coli* BL21(DE3) (200 ng) was incubated with 0.5 μg of purified VPA1263 variants for 5 min at 30°C in DNase assay buffer (20 mM Tris-Cl pH 7.5, 200 mM NaCl, 5% vol/vol glycerol) supplemented with either 2 mM EDTA (a metal chelator) or MgCl_2_. The total volume of the reactions was 20 μL. The reactions were stopped by adding 6.65 μg of Proteinase K and the samples were incubated for 5 min at 55°C. Samples were analyzed by 1.0% agarose-gel electrophoresis. For positive and negative controls, 1 U DNase I (Thermo Fisher Scientific) and 0.5 μg BSA (Sigma) were used, respectively. The experiments were performed twice with similar results. Results from a representative experiment are shown.

### In vivo DNase assays

*E. coli* MG1655 strains containing the indicated pBAD^K^/Myc-His plasmid, either empty or encoding VPA1263^547-869^ (PyS-HNH), VPA1263^547-869/H837A^ (PyS-HNH^HA^), or PoNe^Bc^ (BC3021), were grown overnight in 2xYT media supplemented with kanamycin and 0.4% (wt/vol) glucose. Overnight cultures were washed with 2xYT media and normalized to an OD_600_ of 1.0 in 3 mL of fresh 2xYT supplemented with kanamycin and 0.1% (wt/vol) L-arabinose (to induce protein expression). Cultures were grown for 90 min with agitation (220 rpm) at 37°C before 1.0 OD_600_ units were pelleted. Genomic DNA was isolated from each sample using the EZ spin column genomic DNA kit (Bio Basic) and eluted with 30 μL of ultrapure water (Milli-Q). Equal genomic DNA elution volumes were analyzed by 1.0% (wt/vol) agarose gel electrophoresis. The experiments were performed at least three times with similar results. Results from a representative experiment are shown.

### *Vibrio* growth assays

*Vibrio* growth assays were performed as previously described (21). Briefly, overnight cultures of *V. parahaemolyticus* strains were normalized to OD_600_ = 0.01 in MLB and transferred to 96-well plates (200 μL per well; *n* = 3 technical replicates). The 96-well plates were incubated in a microplate reader (BioTek SYNERGY H1) at 30°C with constant shaking at 205 cpm. OD_600_ reads were acquired every 10 min. The experiments were performed at least three times with similar results. Results from a representative experiment are shown.

### Bacterial competition assays

Bacterial competition assays were performed as previously described (21). Briefly, attacker and prey strains were grown overnight in MLB with the addition of antibiotics when maintenance of plasmids was required. Bacterial cultures were normalized to OD_600_ = 0.5, and mixed at a 4:1 (attacker:prey) ratio. The mixtures were spotted on MLB agar plates and incubated for 4 hours at 30°C. Plates were supplemented with 0.1% (wt/vol) L-arabinose when expression from an arabinose-inducible plasmid in the attacker strain was required. Colony forming units (CFU) of the prey strain were determined by growing the mixtures on selective plates at the 0 and 4-hour timepoints. The experiments were performed at least three times with similar results. Results from a representative experiment are shown.

### Fluorescence microscopy

Assessment of prey DNA morphology during bacterial competition was performed as previously described (22). Briefly, the indicated *V. parahaemolyticus* RIMD 2210633 strains were grown overnight and mixed as described above for the bacterial competition assays. The Δ*vpa1263*-*vti2* prey strain constitutively expressed GFP (from a stable, high copy number plasmid (54)) to distinguish them from attacker cells. Attacker:prey mixtures were spotted on MLB agar plates and incubated at 30°C for 3 hours. Plates were supplemented with 0.1% (wt/vol) L-arabinose when expression from an arabinose-inducible plasmid in the attacker strain was required. Bacteria were then scraped from the plates, washed with M9 media, and incubated at room temperature for 10 min in M9 media containing the Hoechst 33342 (Invitrogen) DNA dye at a final concentration of 1 μg/μL. The cells were then washed and resuspended in 30 μL M9 media. One microliter of bacterial suspension was spotted onto M9 agar (1.5% wt/vol) pads and allowed to dry for 2 min before the pads were placed face-down in 35 mm glass bottom Cellview cell culture dishes. Bacteria were imaged in a Nikon Eclipse Ti2-E inverted motorized microscope equipped with a CFI PLAN apochromat DM 100X oil lambda PH-3 (NA, 1.45) objective lens, a Lumencor SOLA SE II 395 light source, and ET-DAPI (#49028, used to visualize Hoechst signal) and ET-EGFP (#49002, used to visualize GFP signal) filter sets. Images were acquired using a DS-QI2 Mono cooled digital microscope camera (16 MP), and were post-processed using Fiji ImageJ suite. Cells exhibiting a normal DNA morphology or aberrant DNA morphology (i.e., DNA foci or no DNA) were manually counted (> 100 GFP-expressing prey cells per treatment in each experiment). The experiment was repeated three times.

### Protein secretion assays

*V. parahaemolyticus* RIMD 2210633 strains were grown overnight at 30°C in MLB supplemented with kanamycin to maintain plasmids. Bacterial cultures were normalized to OD_600_ = 0.18 in 5 mL MLB supplemented with kanamycin and L-arabinose (0.05% wt/vol) to induce expression from P*bad* promoters. After 4 hours of incubation at 30°C with agitation (220 rpm), 1.0 OD_600_ units of cells were collected for expression fractions (cells). The cell pellets were resuspended in (2X) Tris-Glycine SDS sample buffer (Novex, Life Sciences). For secretion fractions (media), suspension volumes equivalent to 10 OD_600_ units of cells were filtered (0.22 μm), and proteins were precipitated from the media using deoxycholate and trichloroacetic acid (55). Cold acetone was used to wash the protein precipitates twice. Then, protein precipitates were resuspended in 20 μL of 10 mM Tris-HCl pH = 8, followed by the addition of 20 μL of (2X) Tris-Glycine SDS Sample Buffer supplemented with 5% β-mercaptoethanol and 0.5 μL 1 N NaOH to maintain a basic pH. Expression and secretion samples were boiled for 5 min and then resolved on Mini-PROTEAN or Criterion™ TGX Stain-Free™ precast gels (Bio-Rad). For immunoblotting, primary antibodies were used at a 1:1000 concentration. The following antibodies were used: α-Myc antibodies (Santa Cruz Biotechnology, 9E10, mouse mAb; sc-40) and custom-made α-VgrG1 (56). Protein signals were visualized in a Fusion FX6 imaging system (Vilber Lourmat) using enhanced chemiluminescence (ECL) reagents.

### VPA1263 domain and homologs analysis

Domains in VPA1263 (BAC62606.1) were identified using the NCBI conserved domain database (31) or by homology detection and structure prediction using a hidden Markov model (HMM)-HMM comparison, i.e., the HHpred tool (34). Proteins containing a specialized secretion system delivery domain or known secreted toxins bearing homology to the C-terminal HNH nuclease toxin domain of VPA1263 were identified through an iterative search against the UniProt protein sequence database using Jackhmmer (57). Homologs containing diverse secretion-associated domains (e.g., domains found in T6SS or T7SS polymorphic toxins) were selected, and their C-terminal ends were aligned in MEGA X (58) using MUSCLE (59); alignment columns not represented in VPA1263 were removed. Conserved residues were illustrated using the WebLogo 3 server (60). Amino acid numbering was based on the sequence of VPA1263.

## ACKNOWLEDGMENTS

This project received funding from the European Research Council under the European Union’s Horizon 2020 research and innovation program (grant agreement no. 714224), and from the Israel Science Foundation (grant no. 920/17 to D Salomon, and grant no. 1362/21 to D Salomon and E Bosis). CM Fridman was supported by scholarships from the Clore Israel Foundation and from the Manna Center Program in Food Safety and Security at Tel Aviv University, as well as by a scholarship for outstanding doctoral students from the Orthodox community from the Council for Higher Education. The funders had no role in study design, data collection and interpretation, or the decision to submit the work for publication. We thank members of the Salomon lab for helpful discussions and suggestions, and Dan Goldenberg for technical assistance. This work was performed in partial fulfillment of the requirements for a PhD degree for CM Fridman at the Sackler Faculty of Medicine, Tel Aviv University.

## AUTHOR CONTRIBUTIONS

CM Fridman: conceptualization, investigation, methodology, and writing—original draft.

B Jana: conceptualization, investigation, methodology, and writing—review and editing.

R Ben-Yaakov: investigation and methodology.

E Bosis: conceptualization, investigation, methodology, funding acquisition, and writing—review and editing.

D Salomon: conceptualization, supervision, funding acquisition, investigation, methodology, and writing—original draft.

## REFERENCES

1. Jamet A, Nassif X. 2015. New players in the toxin field: polymorphic toxin systems in bacteria. MBio 6:e00285-15.

2. Zhang D, de Souza RF, Anantharaman V, Iyer LM, Aravind L. 2012. Polymorphic toxin systems: Comprehensive characterization of trafficking modes, processing, mechanisms of action, immunity and ecology using comparative genomics. Biol Direct 7:18.

3. Salomon D, Kinch LN, Trudgian DC, Guo X, Klimko JA, Grishin N V., Mirzaei H, Orth K. 2014. Marker for type VI secretion system effectors. Proc Natl Acad Sci 111:9271–9276.

4. Ruhe ZC, Low DA, Hayes CS. 2020. Polymorphic toxins and their immunity proteins: Diversity, evolution, and mechanisms of delivery. Annu Rev Microbiol 74:497–520.

5. Whitney JC, Peterson SB, Kim J, Pazos M, Verster AJ, Radey MC, Kulasekara HD, Ching MQ, Bullen NP, Bryant D, Goo YA, Surette MG, Borenstein E, Vollmer W, Mougous JD. 2017. A broadly distributed toxin family mediates contact-dependent antagonism between gram-positive bacteria. Elife 6:e26938.

6. Aoki SK, Diner EJ, de Roodenbeke C t’Kint, Burgess BR, Poole SJ, Braaten BA, Jones AM, Webb JS, Hayes CS, Cotter PA, Low DA. 2010. A widespread family of polymorphic contact-dependent toxin delivery systems in bacteria. Nature 468:439–442.

7. Pukatzki S, Ma AT, Sturtevant D, Krastins B, Sarracino D, Nelson WC, Heidelberg JF, Mekalanos JJ. 2006. Identification of a conserved bacterial protein secretion system in Vibrio cholerae using the Dictyostelium host model system. Proc Natl Acad Sci 103:1528–1533.

8. Boyer F, Fichant G, Berthod J, Vandenbrouck Y, Attree I. 2009. Dissecting the bacterial type VI secretion system by a genome wide in silico analysis: What can be learned from available microbial genomic resources? BMC Genomics 10:104.

9. Silverman JM, Agnello DM, Zheng H, Andrews BT, Li M, Catalano CE, Gonen T, Mougous JD. 2013. Haemolysin coregulated protein is an exported receptor and chaperone of type VI secretion substrates. Mol Cell 51:584–593.

10. Hachani A, Allsopp LP, Oduko Y, Filloux A. 2014. The VgrG proteins are “à la carte” delivery systems for bacterial type VI effectors. J Biol Chem 289:17872–84.

11. Burkinshaw BJ, Liang X, Wong M, Le ANH, Lam L, Dong TG. 2018. A type VI secretion system effector delivery mechanism dependent on PAAR and a chaperone-co-chaperone complex. Nat Microbiol 3:632–640.

12. Wang J, Brodmann M, Basler M. 2019. Assembly and subcellular localization of bacterial type VI secretion systems. Annu Rev Microbiol 73.

13. Basler M, Pilhofer M, Henderson GP, Jensen GJ, Mekalanos JJ. 2012. Type VI secretion requires a dynamic contractile phage tail-like structure. Nature 483:182–6.

14. Jana B, Salomon D. 2019. Type VI secretion system: a modular toolkit for bacterial dominance. Future Microbiol 14:fmb-2019-0194.

15. Jurėnas D, Journet L. 2021. Activity, delivery, and diversity of Type VI secretion effectors. Mol Microbiol 115:383–394.

16. Unterweger D, Kostiuk B, Ötjengerdes R, Wilton A, Diaz-Satizabal L, Pukatzki S. 2015. Chimeric adaptor proteins translocate diverse type VI secretion system effectors in Vibrio cholerae. EMBO J 34:2198–210.

17. Liang X, Moore R, Wilton M, Wong MJQ, Lam L, Dong TG. 2015. Identification of divergent type VI secretion effectors using a conserved chaperone domain. Proc Natl Acad Sci 112:9106–9111.

18. Bondage DD, Lin J-S, Ma L-S, Kuo C-H, Lai E-M. 2016. VgrG C terminus confers the type VI effector transport specificity and is required for binding with PAAR and adaptor–effector complex. Proc Natl Acad Sci 113:E3931–E3940.

19. Alcoforado Diniz J, Coulthurst SJ. 2015. Intraspecies competition in Serratia marcescens is mediated by type VI-secreted Rhs effectors and a conserved effector-associated accessory protein. J Bacteriol 197:2350–60.

20. Cianfanelli FR, Alcoforado Diniz J, Guo M, De Cesare V, Trost M, Coulthurst SJ. 2016. VgrG and PAAR proteins define distinct versions of a functional type VI secretion system. PLoS Pathog 12:1–27.

21. Dar Y, Jana B, Bosis E, Salomon D. 2021. A binary effector module secreted by a type VI secretion system. EMBO Rep e53981.

22. Jana B, Fridman CM, Bosis E, Salomon D. 2019. A modular effector with a DNase domain and a marker for T6SS substrates. Nat Commun 10:3595.

23. Koskiniemi S, Lamoureux JG, Nikolakakis KC, t’Kint de Roodenbeke C, Kaplan MD, Low DA, Hayes CS. 2013. Rhs proteins from diverse bacteria mediate intercellular competition. Proc Natl Acad Sci U S A 110:7032–7.

24. Miyata ST, Kitaoka M, Brooks TM, McAuley SB, Pukatzki S. 2011. Vibrio cholerae requires the type VI secretion system virulence factor vasx to kill dictyostelium discoideum. Infect Immun 79:2941–2949.

25. Salomon D, Klimko JA, Trudgian DC, Kinch LN, Grishin N V., Mirzaei H, Orth K. 2015. Type VI secretion system toxins horizontally shared between marine bacteria. PLoS Pathog 11:1–20.

26. Ray A, Schwartz N, Souza Santos M, Zhang J, Orth K, Salomon D, de Souza Santos M, Zhang J, Orth K, Salomon D. 2017. Type VI secretion system MIX-effectors carry both antibacterial and anti-eukaryotic activities. EMBO Rep 18:e201744226.

27. Hood RD, Singh P, Hsu FS, Güvener T, Carl MA, Trinidad RRS, Silverman JM, Ohlson BB, Hicks KG, Plemel RL, Li M, Schwarz S, Wang WY, Merz AJ, Goodlett DR, Mougous JD. 2010. A type VI secretion system of Pseudomonas aeruginosa targets a toxin to bacteria. Cell Host Microbe 7:25–37.

28. Russell AB, Singh P, Brittnacher M, Bui NK, Hood RD, Carl MA, Agnello DM, Schwarz S, Goodlett DR, Vollmer W, Mougous JD. 2012. A widespread bacterial type VI secretion effector superfamily identified using a heuristic approach. Cell Host Microbe 11:538–549.

29. Dar Y, Salomon D, Bosis E. 2018. The antibacterial and anti-eukaryotic Type VI secretion system MIX-effector repertoire in Vibrionaceae. Mar Drugs 16:433.

30. Hurley CC, Quirke AM, Reen FJ, Boyd EF. 2006. Four genomic islands that mark post-1995 pandemic Vibrio parahaemolyticus isolates. BMC Genomics 7:104.

31. Marchler-Bauer A, Anderson JB, Derbyshire MK, DeWeese-Scott C, Gonzales NR, Gwadz M, Hao L, He S, Hurwitz DI, Jackson JD, Ke Z, Krylov D, Lanczycki CJ, Liebert CA, Liu C, Lu F, Lu S, Marchler GH, Mullokandov M, Song JS, Thanki N, Yamashita RA, Yin JJ, Zhang D, Bryant SH. 2007. CDD: a conserved domain database for interactive domain family analysis. Nucleic Acids Res 35:D237-40.

32. Buist G, Steen A, Kok J, Kuipers OP. 2008. LysM, a widely distributed protein motif for binding to (peptido)glycans. Mol Microbiol 68:838–847.

33. Michel-Briand Y, Baysse C. 2002. The pyocins of Pseudomonas aeruginosa. Biochimie https://doi.org/10.1016/S0300-9084(02)01422-0.

34. Zimmermann L, Stephens A, Nam S-Z, Rau D, Kübler J, Lozajic M, Gabler F, Söding J, Lupas AN, Alva V. 2018. A completely reimplemented MPI bioinformatics toolkit with a new HHpred server at its core. J Mol Biol 430:2237–2243.

35. Saravanan M, Bujnicki JM, Cymerman IA, Rao DN, Nagaraja V. 2004. Type II restriction endonuclease R.Kpnl is a member of the HNH nuclease superfamily. Nucleic Acids Res 32:6129–6135.

36. Sano Y, Matsui H, Kobayashi M, Kageyama M. 1993. Molecular structures and functions of pyocins S1 and S2 in Pseudomonas aeruginosa. J Bacteriol 175:2907–2916.

37. Chak KF, Kuo WS, Lu FM, James R. 1991. Cloning and characterization of the ColE7 plasmid. J Gen Microbiol 137:91–100.

38. Landthaler M, Lau NC, Shub DA. 2004. Group I intron homing in Bacillus phages SPO1 and SP82: A gene conversion event initiated by a nicking homing endonuclease. J Bacteriol 186.

39. Stoddard BL. 2005. Homing endonuclease structure and function. Q Rev Biophys https://doi.org/10.1017/S0033583505004063.

40. Kala S, Cumby N, Sadowski PD, Hyder BZ, Kanelis V, Davidson AR, Maxwell KL. 2014. HNH proteins are a widespread component of phage DNA packaging machines. Proc Natl Acad Sci U S A 111:6022–7.

41. Poulsen C, Panjikar S, Holton SJ, Wilmanns M, Song Y-H. 2014. WXG100 Protein Superfamily Consists of Three Subfamilies and Exhibits an α-Helical C-Terminal Conserved Residue Pattern. PLoS One 9:e89313.

42. Klein A, Wojdyla JA, Joshi A, Josts I, McCaughey LC, Housden NG, Kaminska R, Byron O, Walker D, Kleanthous C. 2016. Structural and biophysical analysis of nuclease protein antibiotics. Biochem J 473:2799–812.

43. Salomon D, Klimko JA, Orth K. 2014. H-NS regulates the Vibrio parahaemolyticus type VI secretion system 1. Microbiol (United Kingdom) 160:1867–1873.

44. Levin-Zaidman S, Frenkiel-Krispin D, Shimoni E, Sabanay I, Wolf SG, Minsky A. 2000. Ordered intracellular RecA-DNA assemblies: A potential site of in vivo RecA-mediated activities. Proc Natl Acad Sci U S A 97.

45. Atanaskovic I, Sharp C, Press C, Kaminska R, Kleanthous C. 2022. Bacterial Competition Systems Share a Domain Required for Inner Membrane Transport of the Bacteriocin Pyocin G from Pseudomonas aeruginosa. MBio 13:e0339621.

46. Vettiger A, Basler M. 2016. Type VI Secretion System Substrates Are Transferred and Reused among Sister Cells. Cell 167:99-110.e12.

47. Quentin D, Ahmad S, Shanthamoorthy P, Mougous JD, Whitney JC, Raunser S. 2018. Mechanism of loading and translocation of type VI secretion system effector Tse6. Nat Microbiol 3:1142–1152.

48. Ho BT, Fu Y, Dong TG, Mekalanos JJ. 2017. *Vibrio cholerae* type 6 secretion system effector trafficking in target bacterial cells. Proc Natl Acad Sci 201711219.

49. Jana B, Keppel K, Salomon D. 2021. Engineering a customizable antibacterial T6SS-based platform in Vibrio natriegens. EMBO Rep 22:e53681.

50. Makino K, Oshima K, Kurokawa K, Yokoyama K, Uda T, Tagomori K, Iijima Y, Najima M, Nakano M, Yamashita A, Kubota Y, Kimura S, Yasunaga T, Honda T, Shinagawa H, Hattori M, Iida T. 2003. Genome sequence of Vibrio parahaemolyticus: a pathogenic mechanism distinct from that of V cholerae. Lancet 361:743–749.

51. Gibson DG, Young L, Chuang RY, Venter JC, Hutchison CA, Smith HO. 2009. Enzymatic assembly of DNA molecules up to several hundred kilobases. Nat Methods 6:343–345.

52. O’Toole R, Milton DL, Wolf-Watz H. 1996. Chemotactic motility is required for invasion of the host by the fish pathogen Vibrio anguillarum. Mol Microbiol 19:625–637.

53. Salomon D, Gonzalez H, Updegraff BL, Orth K. 2013. Vibrio parahaemolyticus Type VI secretion system 1 Is activated in marine conditions to target bacteria, and is differentially regulated from system 2. PLoS One 8:e61086.

54. Ritchie JM, Rui H, Zhou X, Iida T, Kodoma T, Ito S, Davis BM, Bronson RT, Waldor MK. 2012. Inflammation and Disintegration of Intestinal Villi in an Experimental Model for Vibrio parahaemolyticus-Induced Diarrhea. PLoS Pathog 8:e1002593.

55. Bensadoun A, Weinstein D. 1976. Assay of proteins in the presence of interfering materials. Anal Biochem 70:241–250.

56. Li P, Kinch LN, Ray A, Dalia AB, Cong Q, Nunan LM, Camilli A, Grishin N V, Salomon D, Orth K. 2017. Acute hepatopancreatic necrosis disease-causing Vibrio parahaemolyticus strains maintain an antibacterial type VI secretion system with versatile effector repertoires. Appl Environ Microbiol 83:e00737-17.

57. Potter SC, Luciani A, Eddy SR, Park Y, Lopez R, Finn RD. 2018. HMMER web server: 2018 update. Nucleic Acids Res 46:W200–W204.

58. Kumar S, Stecher G, Li M, Knyaz C, Tamura K. 2018. MEGA X: Molecular evolutionary genetics analysis across computing platforms. Mol Biol Evol 35:1547–1549.

59. Edgar RC. 2004. MUSCLE: multiple sequence alignment with high accuracy and high throughput. Nucleic Acids Res 32:1792–1797.

60. Crooks GE, Hon G, Chandonia J-M, Brenner SE. 2004. WebLogo: a sequence logo generator. Genome Res 14:1188–90.

