## Supplementary Information - Figures and Tables for "A DNase T6SS effector requires its MIX domain for secretion"

[a] Department of Clinical Microbiology and Immunology, Sackler Faculty of Medicine, Tel Aviv University, Tel Aviv, Israel

### Supplementary Figures

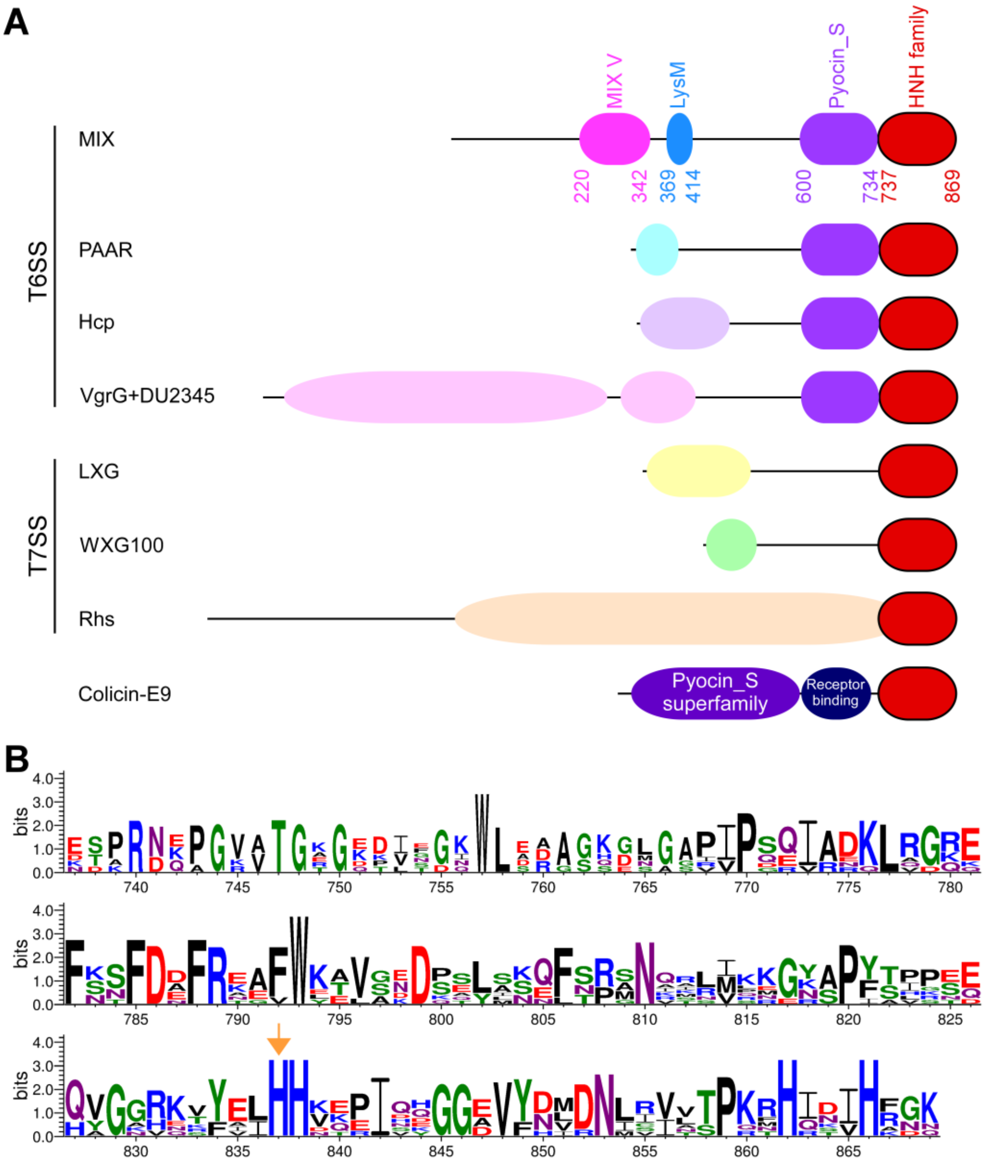


**Supplementary Figure S1. Homology and conservation of VPA1263's C-terminal toxin domain. (A)** Schematic representation of selected homologs of the C-terminal toxin domain in VPA1263. N-terminal domains and predicted secretion systems are denoted on the left. Secretion systems were determined by system-specific domains or neighboring genes. MIX, VPA1263 from *Vibrio parahaemolyticus* RIMD 2210633; PAAR, PQQ40857.1 from *Photorhabdus luminescens* H5; Hcp, SMR99238.1 from *Vibrio mangrovi* CECT 7927; VgrG, EAV0570772.1 from *Salmonella enterica* PNUSAS065855; LXG, EOP65835.1 from *Bacillus cereus* VD118; WXG100, EJP82933.1 from *Bacillus cereus* VD022; Rhs, REH35728.1 from *Kutzneria buriramensis* DSM 45791; Colicin-E9, CAA31104.1 from *Escherichia coli* k-12. **(B)** A conserved motif of the VPA1263 C-terminal HNH family domain, as illustrated using WebLogo 3, based on multiple sequence alignment of the proteins described in **A**. The numbers below denote the positions in VPA1263. An orange arrow denotes the active site Histidine at position 837 of VPA1263.

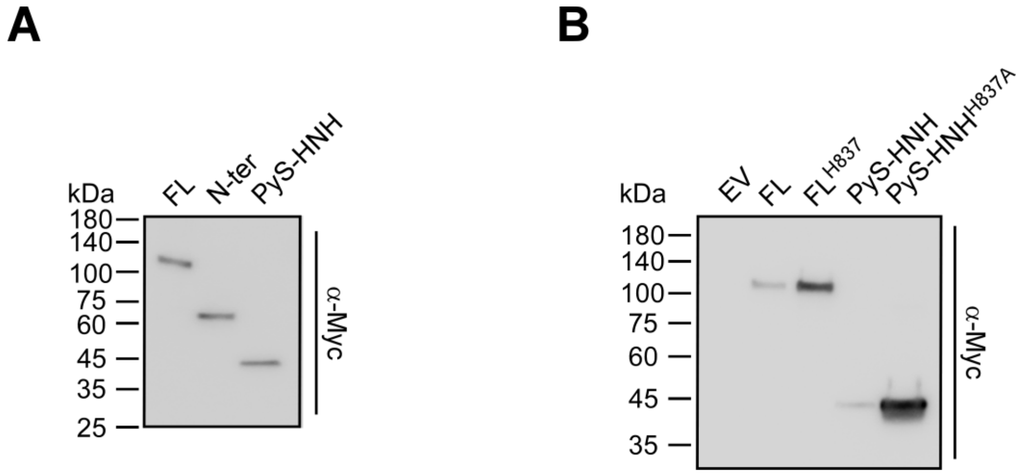


**Supplementary Figure S2. Expression of VPA1263 variants in *E. coli*. (A-B)** Expression of the indicated C-terminal Myc-His-tagged VPA1263 variants from an arabinose-inducible plasmid in *E. coli* MG1655. Proteins were detected by immunoblotting using specific α-Myc antibodies. FL, full-length VPA1263; N-ter, VPA1263 amino acids 1-546; PyS-HNH, VPA1263 amino acids 547-869; EV, empty plasmid; $FL^{HA}$, $VPA1263^{H837A}$; PyS-HNH $^{H837A}$, VPA1263 amino acids 547-869 with a H837A mutation.

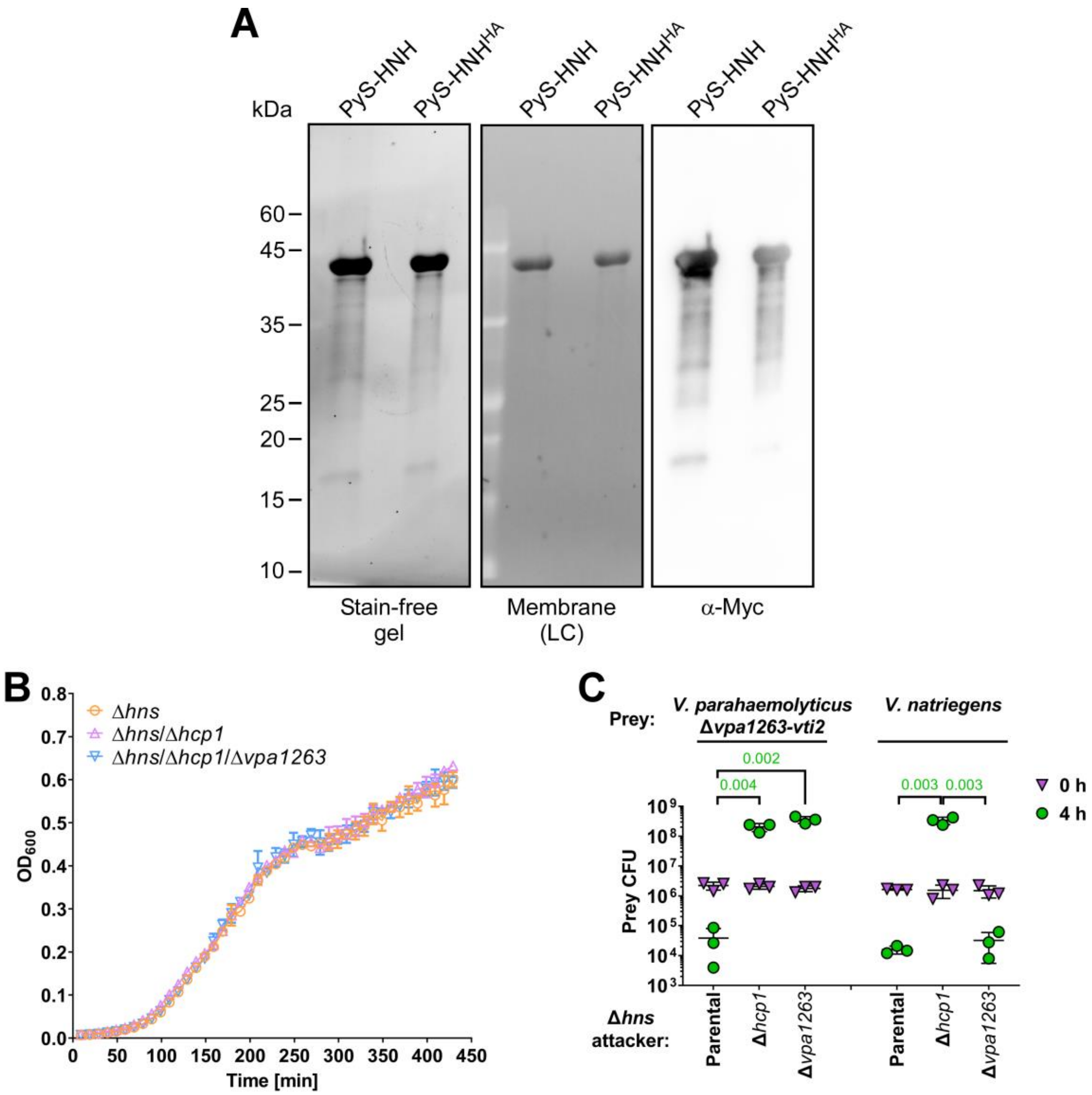


**Supplementary Figure S3. VPA1263 mediates T6SS-dependent killing. (A)** Purification of VPA1263$^{547\text{-}869}$ (PyS-HNH) and VPA1263$^{547\text{-}869/H837}$ (PyS-HNH$^{HA}$) containing a C-terminal Myc-His tag from *E. coli*. Final elution fractions were resolved on TGX stain-free gels (Bio-Rad); total proteins were visualized using the gel's trihalo compounds' fluorescence after UV activation on the gel and after transfer to a nitrocellulose membrane (LC, loading control). VPA1263 variants were detected by immunoblotting with α-Myc antibodies. **(B)** Growth of the indicated *V. parahaemolyticus* RIMD 2210633 derivatives in MLB at 30°C is shown as optical density measurements at 600 nm ($OD_{600}$). Data are shown as the mean ± SD, n=3 technical repeats. The experiment was performed three times with similar results; results of a representative experiment are shown. **(C)** Viability counts of the indicated prey strains before (0 h) and after (4 h) co-incubation with the indicated *V. parahaemolyticus* RIMD 2210633 Δ*hns* derivative attacker strain on MLB agar plates at 30°C. Prey strains contain an empty plasmid that provides a selection marker. Data are shown as the mean ± SD, n = 3 technical replicates. Statistical significance between samples at the 4 h timepoint by an unpaired two-tailed Student's *t*-test is denoted above. The experiment was performed three times with similar results; results of a representative experiment are shown.

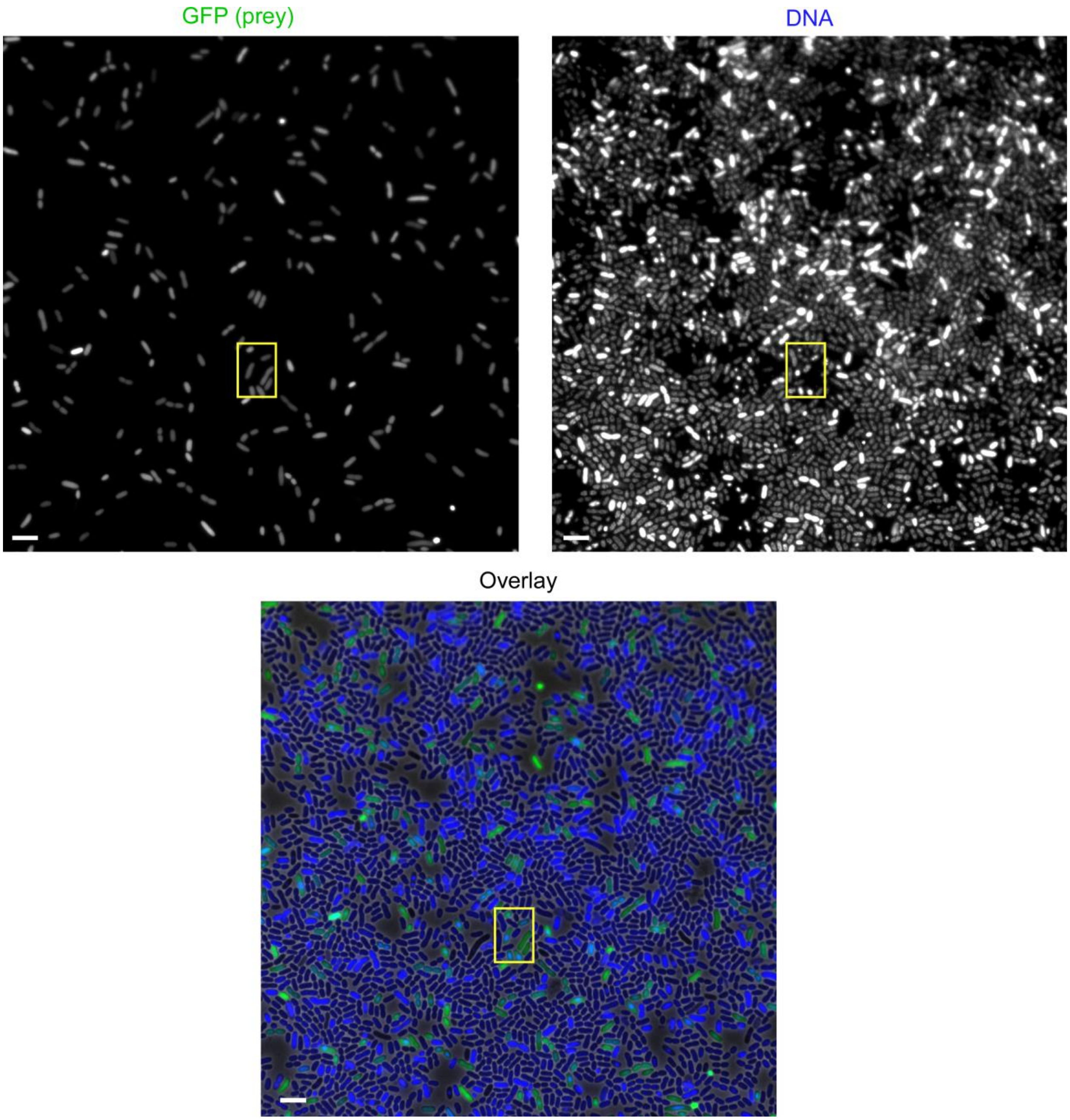


**Supplementary Figure S4. VPA1263-mediated toxicity results in aberrant prey DNA morphology during bacterial competition.** The complete field of view of fluorescence microscope images shown in Figure 2D. The section shown in Figure 2D is denoted with a yellow rectangle. Bar = 4 µm.

### Supplementary Tables

**Supplementary Table S1. A list of bacterial strains used in this study.**

| Strain name | Genotype | Comments | Source |
|---|---|---|---|
| *Vibrio parahaemolyticus* RIMD 2210633 | Wild type | Used for generating deletion strains and as a template for PCR amplification | Obtained from Kim Orth; (1) |
| *Vibrio parahaemolyticus* RIMD 2210633 Δ*hns* | Δ*hns* | A RIMD 2210633 derivative containing an in-frame deletion of *vp1133*; it is used as an attacker in bacterial competition assays, and in secretion assays | (2) |
| *Vibrio parahaemolyticus* RIMD 2210633 Δ*hns*/Δ*hcp1* | Δ*hns*/Δ*hcp1* | A RIMD 2210633 derivative containing an in-frame deletion of *vp1133* and of *vp1393*; it is used as an attacker in bacterial competition assays and in secretion assays | (2) |
| *Vibrio parahaemolyticus* RIMD 2210633 Δ*hns*/Δ*vpa1263* | Δ*hns*/Δ*vpa1263* | A RIMD 2210633 derivative containing an in-frame deletion of *vp1133* and of *vpa1263*; it is used as an attacker in bacterial competition assays | This study |

| Strain name | Genotype | Comments | Source |
|---|---|---|---|
| *Vibrio parahaemolyticus* RIMD 2210633 Δ*vpa1263-vti2* | Δ*vpa1263-vti2* | A RIMD 2210633 derivative containing an in-frame deletion of *vpa1263-vti2*; it is used as a prey in bacterial competition assays | This study |
| *Vibrio parahaemolyticus* POR1 $vp1415^{AAA}$ | Δ*tdhAS*/$vp1415^{AAA}$ | A RIMD 2210633 derivative. It is used as a template for PCR amplifications. Codons for histidines 563-4 in VP1415 were replaced by codons for alanines | (3) |
| *Vibrio natriegens* ATCC 14048 | Wild type | Used as a prey in competition assays | ATCC collection |
| *Escherichia coli* DH5α (λ pir) | K-12 derivative laboratory strain containing λ pir | Used for plasmid construction and maintenance | Obtained from Eric V. Stabb |
| *Escherichia coli* MG1655 | Wild type | Used for protein expression, toxicity assays, and in vivo DNase assays | Lab stocks |
| *Escherichia coli* BL21 (DE3) | Laboratory strain | Used for protein expression | Lab stocks |

**Supplementary Table S2. A list of plasmids used in this study.**

| Plasmid name | Description | Comments | Source |
|---|---|---|---|
| pDM4 | a $Cm^R$ and oriVR6K-containing suicide vector | Used to generate deletions in *Vibrio parahaemolyticus* | (4) |
| pDM4:*vpa1263* | pDM4 containing 1 kb upstream and 1 kb downstream of *vpa1263* in its MCS | Used to delete *vpa1263* in *V. parahaemolyticus* RIMD 2210633 | (5) |
| pDM4:*vpa1263-vti2* | pDM4 containing 1 kb upstream of *vpa1263* and 1 kb downstream of *vti2* in its MCS | Used to delete *vpa1263-vti2* in *V. parahaemolyticus* RIMD 2210633 | (5) |
| pDM4:*hns* | pDM4 containing 1 kb upstream and 1 kb downstream of *vp1133* (*hns*) in its MCS | Used to delete *vp1133* in *V. parahaemolyticus* RIMD 2210633 | (6) |
| $pBAD^K$/Myc-His | pBR322 ori-containing plasmid harboring a $Kan^R$ cassette, *araC*, and an MCS following a P*bad* promoter | Used for arabinose-inducible expression | (7) |
| $pPoNe^{BC}$ | $pBAD^K$/Myc-His plasmid containing the CDS of BC3021 ($PoNe^{BC}$) from *Bacillus cereus* ATCC 14579, in frame with a C-terminal Myc-His tag | Used for arabinose-inducible expression of $PoNe^{BC}$ in *E. coli* | (8) |
| pVPA1263 | $pBAD^K$/Myc-His plasmid containing the CDS of *vpa1263,* in frame with a C- | Used for arabinose-inducible expression of VPA1263 (FL) | This study |

| Plasmid name | Description | Comments | Source |
|---|---|---|---|
| | terminal Myc-His tag | during toxicity assays in *E. coli* | |
| pVPA1263$^{1-546}$ | pBAD$^{K}$/Myc-His plasmid containing the CDS of the N-terminal part of *vpa1263* (amino acids 1-546), in frame with a C-terminal Myc-His tag | Used for arabinose-inducible expression of VPA1263$^{1-546}$ (N-ter) during toxicity assays in *E. coli* and during secretion assays in *Vibrio* | This study |
| pVPA1263$^{547-869}$ | pBAD$^{K}$/Myc-His plasmid containing the CDS of the C-terminal part of *vpa1263* (amino acids 547-869), in frame with a C-terminal Myc-His tag | Used for arabinose-inducible expression of VPA1263$^{547-869}$ (PyS-HNH) during toxicity assays and in vivo DNase assays in *E. coli*, and for protein purification | This study |
| pVPA1263$^{737-869}$ | pBAD$^{K}$/Myc-His plasmid containing the CDS of the C-terminal HNH nuclease domain of VPA1263 (amino acids 737-869), in frame with a C-terminal Myc-His tag | Used for the arabinose-inducible expression of VPA1263$^{737-869}$ (HNH) during toxicity assays in *E. coli* | This study |
| pVPA1263$^{H837A}$ | pBAD$^{K}$/Myc-His plasmid containing the CDS of *vpa1263* carrying a mutation in the active site H837A, in frame with a C- | Used for the arabinose-inducible expression of VPA1263$^{H837A}$ (FL$^{HA}$) during toxicity assays in *E. coli* and during | This study |

| Plasmid name | Description | Comments | Source |
| --- | --- | --- | --- |
| | terminal Myc-His tag | secretion assays in *Vibrio* | |
| pVPA1263$^{\Delta MIX/H837A}$ | pBAD$^{K}$/Myc-His plasmid containing the CDS of *vpa1263* lacking most of the MIX domain (amino acids 226-328) and carrying a mutation in the active site, H837A, in frame with a C-terminal Myc-His tag | Used for the arabinose-inducible expression of VPA1263$^{\Delta MIX/H837A}$ ($\Delta$MIX$^{HA}$)during secretion assays in *Vibrio* | This study |
| pVPA1263$^{547\text{-}869/H837A}$ | pBAD$^{K}$/Myc-His plasmid containing the CDS of the C-terminal part of *vpa1263* (amino acids 547-869) and carrying a mutation in the active site H837A, in frame with a C-terminal Myc-His tag | Used for the arabinose-inducible expression of VPA1263$^{547\text{-}869/H837A}$ (PyS-HNH$^{HA}$) during toxicity assays and in vivo DNase assays in *E. coli*, for protein purifications, and during secretion assays in *Vibrio* | This study |
| pVPA1263$^{G247A/H837A}$ | pBAD$^{K}$/Myc-His plasmid containing the CDS of *vpa1263* carrying a mutation in the MIX domain, G246A, followed by a mutation in the active site, H837A, in frame | Used for the arabinose-inducible expression of VPA1263$^{G246A/H837A}$ (FL$^{GA/HA}$) during secretion assays in *Vibrio* | This study |

| Plasmid name | Description | Comments | Source |
|---|---|---|---|
| | with a C-terminal Myc-His tag | | |
| pVPA1263$^{Y250A/H837A}$ | pBAD$^{K}$/Myc-His plasmid containing the CDS of *vpa1263* carrying a mutation in the MIX domain, Y250A, followed by a mutation in the active site, H837A, in frame with a C-terminal Myc-His tag | Used for the arabinose-inducible expression of VPA1263 $^{Y250A/H837A}$ (FL$^{YA/HA}$) during secretion assays in *Vibrio* | This study |
| pBAD33.1$^{F}$ | pBAD33.1 with a FLAG tag inserted at the 3' end of the MCS | Used for arabinose-inducible expression | (9) |
| pVti2 | pBAD33.1$^{F}$ plasmid containing the CDS of *vti2,* in frame with a C-terminal FLAG tag | Used for the arabinose-inducible expression of Vti2 during toxicity assays in *E. coli*, and during protein purification | This study |
| pGFP | a Spec$^{R}$-containing, high copy number plasmid for the constitutive expression of GFP in vibrios | Used for the constitutive expression of GFP in *V. parahaemolyticus* prey visualized under a fluorescence microscope | (10) |

| Plasmid name | Description | Comments | Source |
|---|---|---|---|
| pCLTR | *E. coli*-yeast-*Vibrio* shuttle vector; mobilizable; it contains $Spec^R$ and $Cm^R$ | Used to selectively grow *V. natriegens* prey on media containing chloramphenicol during bacterial competition | (3) |
| pVPA1263-Vti2 | $pBAD^K$/Myc-His plasmid containing the CDS of *vpa1263* and *vti2*; *vti2* is in frame with a C-terminal Myc-His tag | Used for the arabinose-inducible expression of VPA1263 and Vti2 during bacterial competition assays | (3) |
| $pVPA1263^{G247A}$-Vti2 | pVPA1263-Vti2 in which the codon for glycine 247 in *vpa1263* was replaced with a codon for alanine | Used for the arabinose-inducible expression of $VPA1263^{G247A}$ and Vti2 during bacterial competition assays | This study |
| $pVPA1263^{H837A}$-Vti2 | pVPA1263-Vti2 in which the codon for histidine 837 in *vpa1263* was replaced with a codon for alanine | Used for the arabinose-inducible expression of $VPA1263^{H837A}$ and Vti2 during bacterial competition assays | This study |

### Supplementary References


1. Makino K, Oshima K, Kurokawa K, Yokoyama K, Uda T, Tagomori K, Iijima Y, Najima M, Nakano M, Yamashita A, Kubota Y, Kimura S, Yasunaga T, Honda T, Shinagawa H, Hattori M, Iida T. 2003. Genome sequence of Vibrio parahaemolyticus: a pathogenic mechanism distinct from that of V cholerae. Lancet 361:743–749.

2. Dar Y, Jana B, Bosis E, Salomon D. 2021. A binary effector module secreted by a type VI secretion system. EMBO Rep e53981.

3. Jana B, Keppel K, Salomon D. 2021. Engineering a customizable antibacterial T6SS-based platform in Vibrio natriegens. EMBO Rep 22:e53681.

4. O'Toole R, Milton DL, Wolf-Watz H. 1996. Chemotactic motility is required for invasion of the host by the fish pathogen Vibrio anguillarum. Mol Microbiol 19:625–637.

5. Salomon D, Kinch LN, Trudgian DC, Guo X, Klimko JA, Grishin N V., Mirzaei H, Orth K. 2014. Marker for type VI secretion system effectors. Proc Natl Acad Sci 111:9271–9276.

6. Salomon D, Klimko JA, Orth K. 2014. H-NS regulates the Vibrio parahaemolyticus type VI secretion system 1. Microbiol (United Kingdom) 160:1867–1873.

7. Salomon D, Gonzalez H, Updegraff BL, Orth K. 2013. Vibrio parahaemolyticus Type VI secretion system 1 Is activated in marine conditions to target bacteria, and is differentially regulated from system 2. PLoS One 8:e61086.

8. Jana B, Fridman CM, Bosis E, Salomon D. 2019. A modular effector with a DNase domain and a marker for T6SS substrates. Nat Commun 10:3595.

9. Fridman CM, Keppel K, Gerlic M, Bosis E, Salomon D. 2020. A comparative genomics methodology reveals a widespread family of membrane-disrupting T6SS effectors. Nat Commun 11:1085.

10. Ritchie JM, Rui H, Zhou X, Iida T, Kodoma T, Ito S, Davis BM, Bronson RT, Waldor MK. 2012. Inflammation and Disintegration of Intestinal Villi in an Experimental Model for Vibrio parahaemolyticus-Induced Diarrhea. PLoS Pathog 8:e1002593.
